# Current-use and legacy contaminants evidence dissolved organic matter transfer and dynamics across a fractured-rock groundwater recharge area

**DOI:** 10.1101/2024.06.03.596186

**Authors:** Christian Zerfaß, Robert Lehmann, Nico Ueberschaar, Kai Uwe Totsche, Georg Pohnert

## Abstract

Anthropogenic activities cause the release of vast amounts of contaminants into the environment which eventually reach even groundwater resources. With usually sparse regulatory monitoring of limited priority compounds, the large spectrum of contaminants, and the intricacies of intra- and inter-annual contaminant dynamics, such as the emergence and mobilisation of contaminants, are easily overlooked. Utilizing an 6-year record of untargeted LC-MS assessment of dissolved organic, we report the detection and tracing of selected environmental chemicals in the Hainich Critical Zone Exploratory (central Germany), representing a model groundwater flow system under different land use. The insect repellent DEET (*N,N*-diethyl-*m*-toluamide) and the coniferous resin acid 7-ODAA (7-oxodehydroabietic acids) show phases of seasonal dynamics in line with their expected periods of release. The legacy herbicides simazine, the triazine transformation product hydroxypropazine, and the flame retardant/plasticiser TPP (triphenyl phosphate) occurred episodically at various locations in the fractured sedimentary bedrock. Within the period of monitoring, extreme weather events (i.e., the severe 2018 drought) and extreme subsurface responses (i.e., 5-year groundwater highstand 2018) likely contributed to long-term organic matter dynamics, potentially causing re-emergence of legacy agrochemicals. This investigation points to the persistence and mobilisation of anthropogenic contaminants, and highlights the importance of long-term combined untargeted and targeted analysis of the groundwater dissolved organic matter for understanding subsurface ecosystems processes. The results add a note of caution for regulatory monitoring since also legacy contaminant levels may considerably vary over time.

**Highlights:**

- 6 years of monitoring with non-target LC-MS screenings; 4-weekly samplings
- 5 current-use and legacy compounds that evidence dissolved organic matter transfer
- Compounds comprise regularly targeted and non-monitored features in groundwater
- Pronounced phases of non-detects help to further investigate surface-subsurface coupling
- This illustrates the need for spatiotemporally highly resolved non-target monitoring

## 1. Introduction

Groundwater is crucial for mankind as a source of clean water for drinking, agriculture, and other industrial and private uses. The United Nations in their recent World Water Development Report reiterate their call for frequent and long-term monitoring of groundwater resources, to reveal emergences and consequences of contaminations and other adverse impacts (United Nations, 2022). In acknowledgment of the importance of groundwater for processes in the life-sustaining “Critical Zone” (CZ), multidisciplinary investigations have turned their attention to groundwater to understand CZ processes (Grant and Dietrich, 2017; Singha and Navarre-Sitchler, 2022). In this context, dissolved organic matter (DOM) is a key investigative target, as it is both a carrier of carbon and a bioavailable energy source for microorganisms from the surface’s primary production. But it also carries environmental contaminants and persistent chemicals that cause long-term problems. These contaminants can serve for the assessment of ecosystem interconnection and functioning due to their marker properties indicating biogeochemical transformations, migration paths and matter flows (Kaiser and Kalbitz, 2012; Lehmann and Totsche, 2020; Li et al., 2017; Schmidt et al., 2011; Zhang et al., 2018).

For instance, pesticide monitoring, being basically regulated in the EU with the Water Framework Directive (WFD; 2000/60/EC), the Groundwater Directive (GWD; 2006/118/EC), Drinking water Directive (DWD; EU 2020/2184), the Plant Protection Products Regulation (PPP; 1107/2009), and differing national implementations, usually involves only very few measurements per year, even though the monitoring frequency is based on the volume of water distributed or produced within a supply zone (DWD). Quality monitoring is also based on a set of known compounds (e.g. 45 priority substances of EU Directive 2013/39/EC “on Priority Substances in the Field of Water Policy”; “contaminants of emerging concern (CEC)” (Zhang et al., 2021; Merel and Snyder, 2016) by targeted assays, and determine the emergence and/or dynamics of the compounds over time. Beyond that, “unregulated” substances are scarcely, or not monitored at all (Lapworth et al., 2019).

Untargeted (also called: non-target) acquisitions using liquid chromatography coupled to mass spectrometry (LC-MS) are increasingly used to monitor vast amounts of compound features (characterised by retention time and mass-per-charge) in aqueous samples. However, also these comprehensive datasets are often only screened for specific patterns of pre-defined and -validated compound sets, as explorative compound elucidation is challenged by the majority of DOM-signals being poorly defined (Hollender et al., 2017; Lai et al., 2022; Moschet et al., 2017), while for other compounds the environmental relevance may be insufficiently understood (Kiefer et al., 2019). Once a compound is identified, untargeted monitoring datasets can be queried for time-traces of compound dynamics (Hollender et al., 2017). While the non-target approach has successfully been applied for surface water bodies, monitoring data for DOM in groundwater systems remains particularly sparse (Zerfaß et al., 2022).

In this study we explored the long-term record from untargeted DOM monitoring bz LC-MS (Zerfaß et al., 2022) in the Hainich Critical Zone Exploratory (Hainich CZE) for the detection of recurring contaminants in groundwater from a recharge area in central Germany (Küsel et al., 2016). The identified, recurring compounds exhibiting considerable dynamics during a 5-year period of 4-weekly groundwater samplings are the insect repellent DEET (*N,N*-diethyl-*m*-toluamide), the conifer resin acid derivative 7-ODAA (7-oxodehydroabietic acid), the legacy triazine herbicide simazine, the legacy triazine degradation product hydroxypropazine, and the flame retardant/plasticiser TPP (triphenyl phosphate). All compounds were identified from non-targeted LC-MS data sets, verified by reference standards, and monitored over time by re-analysis of the parent non-targeted data set.

The Hainich CZE has been established as monitoring site by the collaborative research centre ‘AquaDiva’ (Küsel et al., 2016; Lehmann and Totsche, 2020) to understand the emergence and fate of “surface signals” and their passage from above-to below-ground. Continuously, information on dissolved organic matter (DOM) has been assessed by liquid chromatography coupled to mass spectrometry (LC-MS) since autumn 2014, and MS1 profile data, interpreted by multivariate statistical methods, has demonstrated that DOM patterns vary most particularly in response to water flows through aquifers (Zerfaß et al., 2022). To identify the compounds behind the MS1-profiles, and to unravel their dynamics over time, we have engaged in compound elucidation by tandem-MS (MS2, data-dependent acquisition) and confirmation of signals by pure reference substances. We demonstrate the emergence of all five compounds mentioned above, both in shallow perched groundwater (7 m depth) and deeper phreatic groundwater (∼90 m). While DEET and 7-ODAA show phases of seasonal dynamics in correspondence to their release pattern all compounds exhibit episodically increased intensities. The anomalous high occurrences in 2021 likely resulted from combined drought and recharge events, eventually enforced by a strong mobilisation triggered by the extreme drought in 2018. These dynamics suggest that intraseasonal fast flow modes can translocate organic matter across the shallow bedrock flow system, while soil-retained (legacy) compounds suffer mobilisation under extreme (combined) events. Taken together, the dynamics of concurrent and legacy DOM components further reveal how flow patterns and surface events shape the quality dynamics across groundwater flow systems.

## 2. Materials and Methods

### 2.1 Sampling site and methodology

In the Hanich Critical Zone Exploratory (Hainich CZE, Figure 1), a hillslope transect of monitoring wells access a groundwater flow system in fractured bedrock of Triassic limestone-mudstone alternations with final (or sampling) depths between 7 and 88 m below ground level (Kohlhepp et al., 2017; Küsel et al., 2016; Lehmann and Totsche, 2020). The model catchment in temperate climate comprise mixed land use with unmanaged woodland (national park), and mostly deciduous forest in low-mountain ridge areas, pasture at midslope positions and cropland further downslope (Figure 1). The methodology for environmental monitoring and sampling (Lehmann and Totsche, 2020), sample processing, chromatography and MS1-assessment (Zerfaß et al., 2022) was documented recently. In brief, filtered (1 µm) groundwater samples were subjected to solid phase extraction (SPE) on Strata-X 33 µm polymeric reversed phase cartridges (500 mg, Phenomenex #8B-S100-HCH, Germany) and eluted with acetonitril/methanol 1:1. Upon evaporation of the solvent, samples were reconditioned in methanol/THF 1:1. LC-MS was carried out on a Dionex UltiMate 3000 chromatography system coupled to a Q-Exactive Plus orbitrap mass spectrometer (Thermo Fisher Scientific, Germany). Reversed phase chromatography (C18) was employed with a gradient from 100 % aqueous solvent A (H_2_O, 2 % acetonitrile, 0.1 % formic acid) to 100 % solvent B (acetonitrile). In later sampling campaigns, we collected LC-MS data for both MS1 (default), and MS2 by data-dependent acquisition (dda, TOP-5) on mixed samples: Per each sampling campaign, samples from up to eight monitoring wells were mixed.

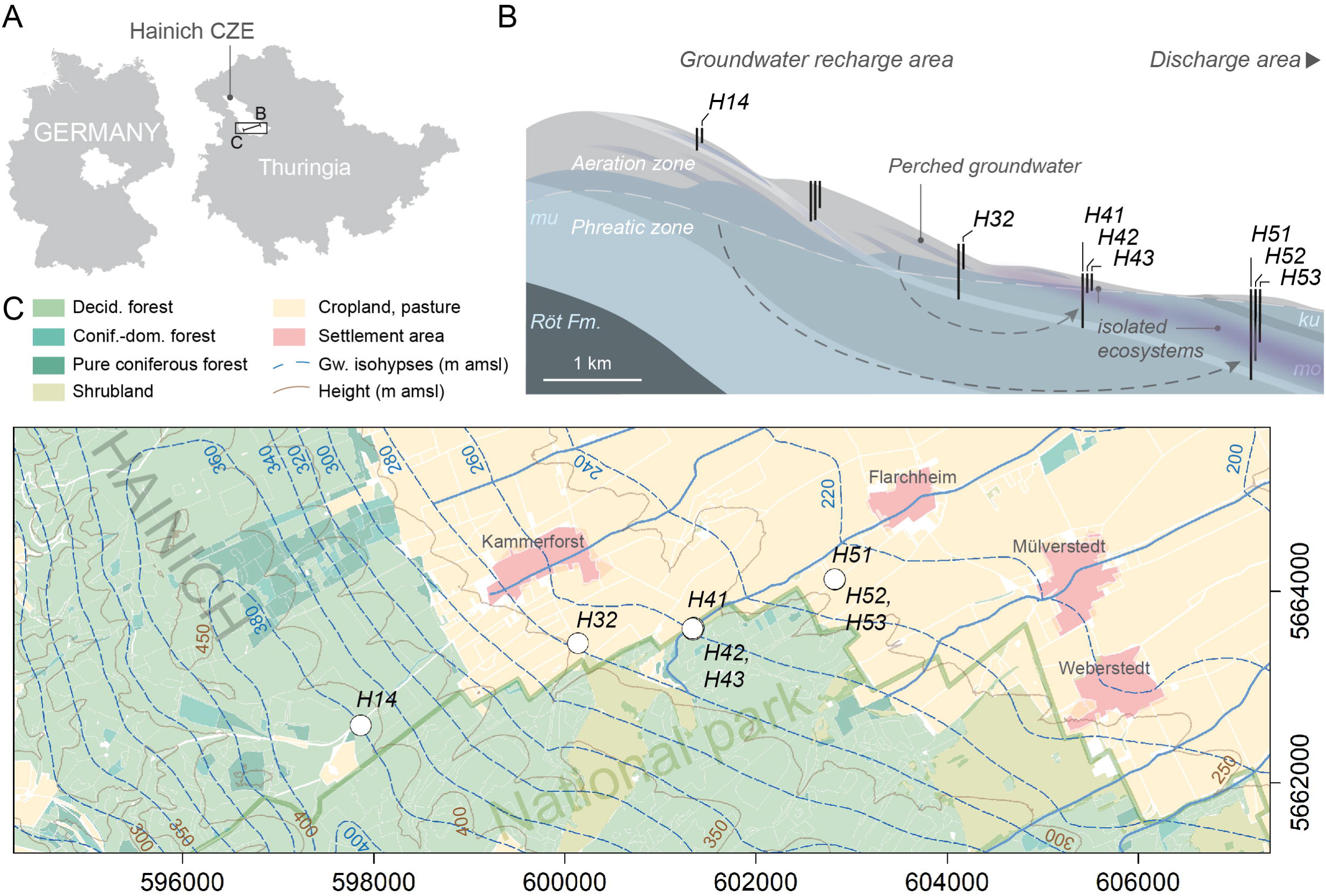

### 2.2 MS-measurements

The MS1 data per sampling campaign was converted by ProteoWizard to the mzXML format (Chambers et al., 2012) and analysed in R (R Core Team, 2022) via the Bioconductor (Huber et al., 2015) package XCMS (Benton et al., 2010; Smith et al., 2006; Tautenhahn et al., 2008) as described previously (Zerfaß et al., 2022). For MS2, a mixed pooled sample per each sampling campaign was prepared, and MS2-data was acquired in dda-mode by which regular MS1-spectra (resolution 70,000 m/Δm, mz-range 100-1,500) were followed by up to five MS2 fragmentation spectra for the five highest signals (resolution 17,500 m/Δm, isolation window 0.2 mz, normalised collision energy (NCE) of 30, dynamic exclusion of 10 s for previously measured parent masses). Further settings were: AGC target 3 x 10^6^, spray voltage 3.0 kV (positive mode) and 2.5 kV (negative mode), capillary temperature 360 °C; positive and negative acquisition in separate runs. MS2 data files were converted to mzXML and split with R to associate relevant MS2 spectra with their preceding MS1 spectra (including isotope candidates), both saved as text-files. These spectra were then analysed in the software SIRIUS (Dührkop et al., 2019, 2015) to retrieve sum formula and structure suggestions. MS1 data analysed in this manuscript is made available via the Metabolights repository (Haug et al., 2019) entries MTBLS3450 (Zerfaß et al., 2021) and MTBLS8433 (Zerfaß and Pohnert, 2024a), and MS2 data in entry MTBLS3533 (Zerfaß and Pohnert, 2024b).

### 2.3 Data processing

To filter the MS2 data for relevant signals, MS1 data was processed to obtain compound features that correspond with MS2 parent masses (mz-ranges) and retention times. Furthermore, only features were retained that in the MS1 data had an Anova false discovery rate (FDR) < 0.01 for being significantly variable between the sampling wells in this campaign. This filtering was applied to remove background signals that may have triggered MS2 acquisitions.

The retained MS1-feature- and associated MS2-information was reviewed manually to identify convincing matches to compound reference databases suggested by the SIRIUS software. For promising propositions with appropriate MS2 spectral quality, reference compounds were obtained for confirmatory runs on the LC-MS in our laboratory. For substance confirmation, DEET (Thermo Fisher Scientific (98 %, #114571000), simazine (Merck, #32059-250MG), hydroxypropazine (LGCstandards, #DRE-C16445000) and triphenyl phosphate (Merck, #442829) were used as delivered. 7-ODAA was synthesised from abietic acid (Merck, #00010-25G) based on a previously published two-step oxidation (Nilsson et al., 2008). Compounds were dissolved and diluted in Methanol:THF 1:1 for LC-MS assessment.

Upon identification of a compound, time traces of MS1-signals were retrieved for eight wells that probe both the shallow perched (H14, H32), and phreatic groundwater (H41-H43, H51-H53) (Lehmann and Totsche, 2020). As features from MS1 in LC-MS may at times be poorly defined from primary analysis of untargeted data (i.e., large retention time or mass-per-charge ranges), XCMS (Benton et al., 2010; Smith et al., 2006; Tautenhahn et al., 2008) was used to prepare extracted ion chromatograms (EICs) at the target mass-per-charge value of the respective compound ions ([M+H]^+^, mz +/- 10 ppm), and chromatographic peaks were re-integrated from the EICs for deriving quantitative intensity time traces.

## 3. Results and Discussion

Tandem-MS analysis by data dependent acquisition (DDA) detected DEET, 7-ODAA, simazine, hydroxypropazine, and TPP in samples taken in spring 2021, and pure reference compound injections confirmed the compound identities (Supplementary Figures S1-S5 and Supplementary Tables S1-S10).

DEET (PubChem, NCBI, 4284) (Figure S1, Tables S1-S2) is approved in the European Union as biocide for repelling or attracting pests as stated in its entry in the European REACH database (European Chemicals Agency (ECHA), 2022). Its most common use is as insect repellent spray for skin application (Aronson et al., 2012; Merel and Snyder, 2016). DEET occurrence in groundwater in Europe is widespread (Loos et al., 2010) and it has been found in other surface waters and precipitation (Costanzo et al., 2007; Merel and Snyder, 2016; Quednow and Püttmann, 2009; Stalder et al., 2022; Zhang et al., 2021). As a volatilised compound that has been detected in precipitation, atmospheric dispersion is a likely mode of translocation.

7-ODAA (PubChem, NCBI, #29212) (Figure S2, Tables S3-S4) is a derivative of the abietan resin acid abietic acic (ABA) or dehydroabietic acid (DbA), all of which are produced specifically in coniferous trees (Goels et al., 2022; Otto and Wilde, 2001; Patyra et al., 2022; Phillips and Croteau, 1999). 7- ODAA occurs as an intermediate in microbial aerobic degradation of abietan (Martin et al., 1999) and as (photo)degradation product of resin acids in water phases (Corin et al., 2000). Abietan derivatives including 7-ODAA are also are released as volatiles from the burning of coniferous wood (Oros and Simoneit, 2001; Simoneit et al., 1993) and thus can atmospherically disperse. In fact, both 7-ODAA, and DbA have previously been found in aerosol particles across Europe, with usually a winter elevation (Oliveira et al., 2007), pointing to heating with wood-burning ovens followed by atmospheric dispersion of 7-ODAA as transport route. In further support, a study of atmospheric PM_2.5_ aerosols (particulate matter with a particle diameter of 2.5 μm or less) in the German city Augsburg (∼300 km south of the Hainich site) over the course of two years revealed that the concentrations of abietan and a methyl-ester of DbA in air increase during winter (Schnelle-Kreis et al., 2007). Far-ranging transport can further be inferred from detection of DbA in air particulates over the Pacific (Simoneit, 2004) and Atlantic (Simoneit and Elias, 2000) ocean.

Simazine (PubChem, NCBI, 5216) (Figure S3, Tables S5-S6) is a legacy groundwater contaminant. A formerly widely used triazine herbicide, its market authorisation in the European Union was withdrawn in 2004 (European Commission, 2004), yet it has been widely detected as groundwater and river contaminant until very recently (Kim et al., 2022; Loos et al., 2010; Robinson et al., 2022). Notably, simazine, alongside other herbicides, has been detected in non-agricultural and remote sites (Wang et al., 2018). The triazines simazine and atrazine have been found in air samples in a Czech background site in 2012/2013, and the article reviews (in the SI) preceding reports of even higher atmospheric abundance from before 2004 when both herbicides were in active agricultural use (Degrendele et al., 2016). This suggests far-ranging transport by atmospheric drift, and correspondingly, a 2000 review presented evidence for simazine in rainwater (Dubus et al., 2000).

Hydroxypropazine (PubChem, NCBI, #135461611) (Figure S4, Tables S7-S8), is a degradation / transformation product of the triazine herbicides prometon, prometryn, and propazine (Schollée et al., 2017; Schollée, Jennifer et al., 2020) (referred to as propazine-2-hydroxy in the cited references). Both prometryn and propazine have been banned as herbicides in the EU prior to 2003 (European Commission, 2010, 2002). Prometon is not listed in the European Union Pesticide Database (European Union, 2023), and no regulatory act on either licensing, or subsequently withdrawing prometon for/from market authorisation could be identified – it has thus conceivably not been used in the EU as a herbicide. Hydroxypropazine can thus be assumed as legacy contaminant transformation product, and previous recent studies detected it still in surface waters (Anagnostopoulou et al., 2022).

TPP (PubChem, NCBI, 8289) (Figure S5, Tables S9-S10) is a plasticiser and lubricant / hydraulic fluid additive. It has previously been detected in river water (Robinson et al., 2022; Van Der Veen and De Boer, 2012), urban stormwater runoff (Masoner et al., 2019), soil (Van Der Veen and De Boer, 2012), indoor air and dust (Van Der Veen and De Boer, 2012), outdoor air and vehicle exhausts (Fabiańska et al., 2019). Transport into the environment through atmospheric particles and leachates is thus conceivable. For groundwater, TPP has been reported in multiple samples of spring-water stemming from a karstified aquifer in Croatia (Lukač Reberski et al., 2023). In a groundwater assessment in Jew Jersey (US), plasticisers – to which TPP belongs – were found as common contaminant, though TPP itself in that study was only detected at a low frequency of one of 58 samples (Stiles et al., 2008). In comparison, (Barnes et al., 2008) also found a fire retardant (tri(2-chloroethyl) phosphate) to occur very frequently in U.S. groundwater.

The intensity time traces for each identified compound were extracted from the long-term monitoring dataset based on the exact monoisotopic mz +/- 10 ppm for the protonated ions, and time traces are reported in Figures 2-6. Sampling from H14, the shallowest well, was sparse between September 2015 – December 2018 as the groundwater body did not yield sufficient water to support LC-MS assessments by our analytical protocol. When sampled, intensities were strongly fluctuating, likely due to fast surface- and soil-fed inputs of organic compounds as indicated by strongly fluctuating water levels and sum parameters (Lehmann and Totsche, 2020).

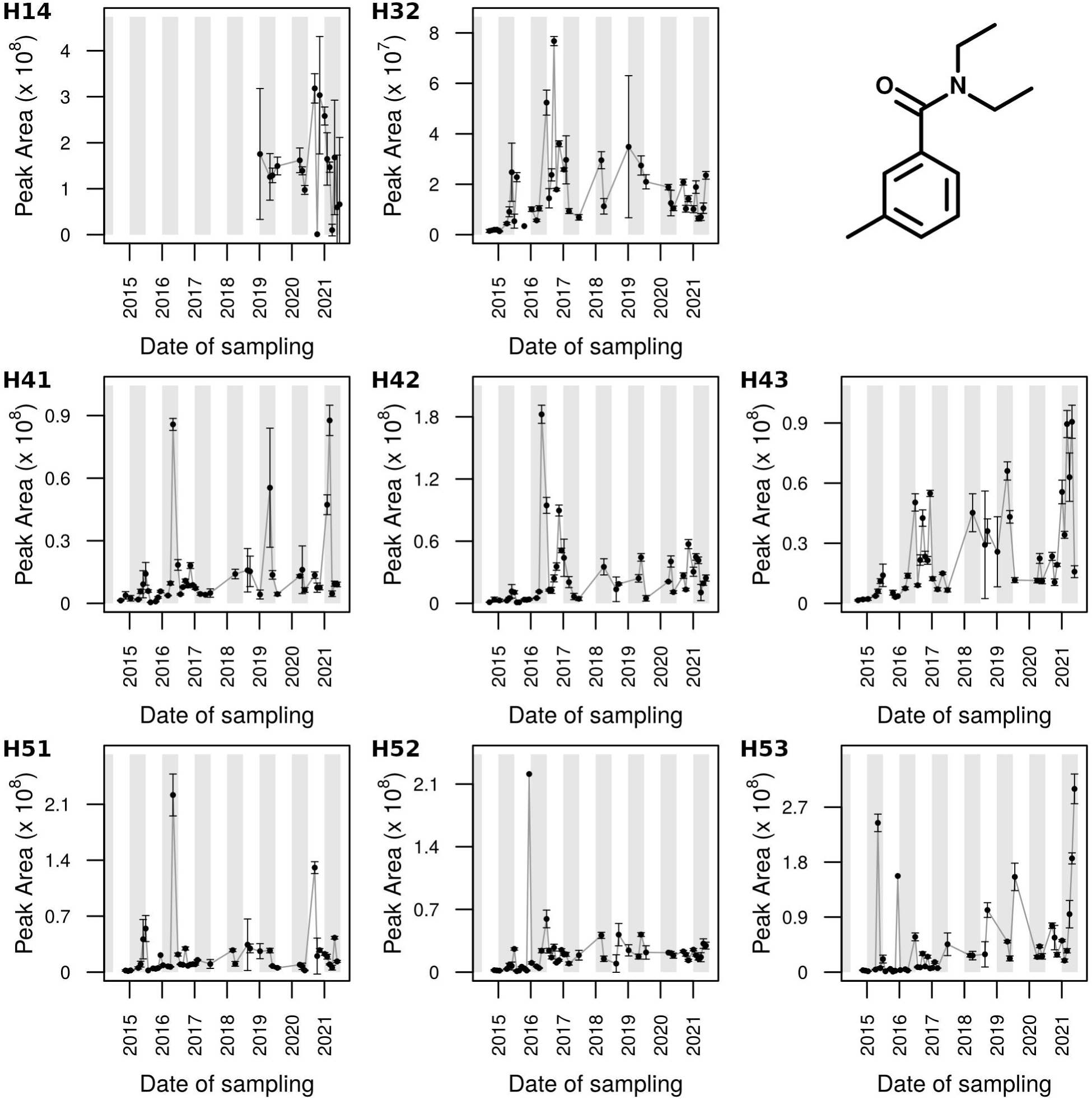

### 3.1 DEET: seasonal diffuse deposition and groundwater loading

In the period 2015-2017, DEET emerged at higher intensities during the (water level) recession periods (Figure 2). This was particularly pronounced in the moderately deeper perched groundwater (H32, 21 m) and the shallow (H42, H43) to deep (H41) phreatic groundwater, where observations over multiple campaigns evidenced a multi-month increase and subsequent decrease. Similar patterns occurred in the phreatic groundwater in discharge direction (H52, H53), where though the high-intensity periods of DEET were much shorter, or only exhibited single-campaign elevations. However, this annual emergence corresponds to the seasonal occurrence in uphill wells. Together with low expected half-life in soil/sediments (<<150 d, see (Weeks et al., 2012)), dynamics point to annual, soil-fed and cross-compartment fluxes. The higher intensities during seasonal highstands/recession (compare Figure 7) periods suggests that DEET is entering the shallow fractured aquifers with infiltration-recharge, after deposition during summer (Knepper, 2004). In accordance, previous studies found higher DEET concentrations in groundwater in summer than in winter (Merel and Snyder, 2016), (Quednow and Püttmann, 2009; Stalder et al., 2022). Furthermore, DEET was reported as being present as water-phase contaminant throughout the year (Quednow and Püttmann, 2009), which reconciles with our detection of DEET in different parts of the year.

Interestingly, after the summer peak in 2016, in several wells the DEET concentration became more fluctuating (e.g., H32, H43) or even demonstrated a pronounced increase to form a defined peak signal towards late autumn 2016 (H41, H42). This also coincided with multiple recharge events (including extreme precipitation, Figure 7 and (Lehmann and Totsche, 2020)) that potentially have caused multiple phases of fast DEET translocation.

In the period from 2017-2020, we acquired less datapoints, though the data consolidates the variable DEET signal intensity over the course of monitoring. Extreme precipitation events in 2017 created a groundwater surplus that protected the groundwater bodies from reaching significant lowstands in response to the severe 2018 summer drought (Peters et al., 2020), whereas DEET fluctuated less strong. In 2021, an interesting signal increase in the first half of the year, starting in the early winter months in some wells, was observed in the phreatic wells H41, H43, and H53 only, that are characterized by contrasting depths and surface-connection. Joint anomalous DEET occurrence and non-detection in other wells during consecutive campaigns, point to more differentiated and always transient flow patterns (Lehmann and Totsche, 2020) or/and retention and transformation within the shallow flow system.

### 3.2 7-ODAA: diffuse deposition or biomarker

In the period 2015-2017, the intensity of 7-ODAA (Figure 3) peaked in winter, with a subsequent decrease towards summer, in the moderately deeper perched (H32) and both shallow and deep phreatic groundwater (H41, H42, H51 and H53). Notably, however, the downward trend was temporarily broken in spring 2015 in wells H32, H41, H42, H51, and H53, likely caused by an episodic recharge event of extreme cumulative precipitation (Lehmann and Totsche, 2020). That highlights the direct impact of atmospheric forcing, or extreme surface conditions for shaping the DOM dynamics in shallow groundwater. Thereafter, the seasonal trend resumed its downward slope. The seasonal pattern corresponds well with the release of 7-ODAA from the burning of coniferous wood (Oros and Simoneit, 2001; Simoneit et al., 1993) in wood-burning ovens during the winter heating period, and correlates with the observation of 7-ODAA in aerosols at higher abundance in winter (Oliveira et al., 2007). While 7-ODAA can form via degradation of coniferous tree tissue (Corin et al., 2000; Martin et al., 1999), our study site is mostly deciduous forest or non-forest surface are (Figure 1 and (Kohlhepp et al., 2017)), and as such the atmospheric input is likely the relevant mechanism. Together with the seasonal dynamics of DEET described in the previous section, the dynamics of 7-ODAA also point to fast seasonal translocation of an atmospherically dispersed compound to the groundwater mobile organic matter pool.

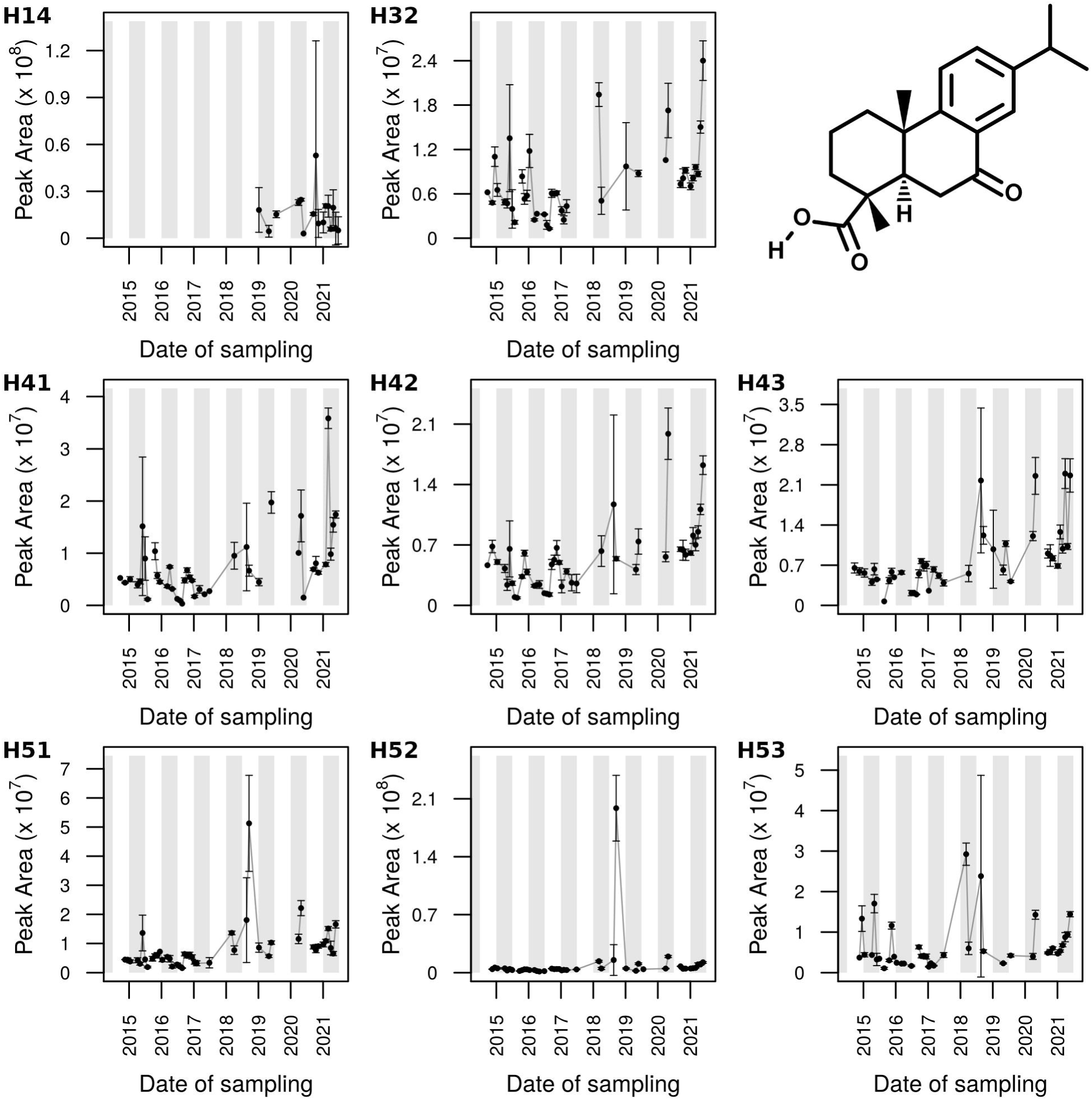

In the first half of 2021, the perched groundwater of well H32 along with phreatic groundwater in wells H41-H43 and H53 showed an increase of intensity for 7-ODAA, which was only weakly indicated in H51 and H52, similar as to the observation for DEET above, though spread across more sampling wells. The wider emergence of this 7-ODAA increase may be explained by the higher hydrophobicity of 7-ODAA (PubChem, NCBI, #29212) compared to DEET (PubChem, NCBI, 4284), so that retention in soil may be stronger – DEET, in fact, has been classed as rather mobile with limited retention in soil (Weeks et al., 2012). Assuming for both a mode of release to the environment via atmospheric dispersion (as argued in the preceding paragraphs), it can be assumed that both compounds may enter the underground fairly ubiquitously, and can thus arrive in different sampling locations. Lack of increased intensities (i.e. H51 and H52, in 2021) for both DEET and 7-ODAA, together with the spring 2015 disruption of seasonal trend in only some wells, collectively highlights the variability of subsurface flow patterns, and of subsurface nutritional supply.

### 3.3 Simazine: a legacy contaminant with intricate dynamics in aeration and phreatic zone wells

The triazine herbicide simazine, which in the EU had its market authorisation withdrawn in 2004 thus over a decade before our first sampling, showed intricate dynamics both in the shallow perched (H14 and H32), and the deeper phreatic zone groundwater (H41 and H51, Figure 4). Its legacy is due to adsorption of the moderately polar pesticide to soil organic matter, resulting in retardation times in vadose zones >10 years (Kim et al., 2022) or even decades (Nanusha et al., 2023), whereas its degradation products can show even longer times (Kim et al., 2022). The absence of simazine in wells H42/H43 and H52/H53, except for intermittent datapoints that are insufficient to infer any dynamic, support earlier findings that characterized these parts of the flow domain as less surface-connected (isolated subsurface ecosystems, see (Lehmann and Totsche, 2020).

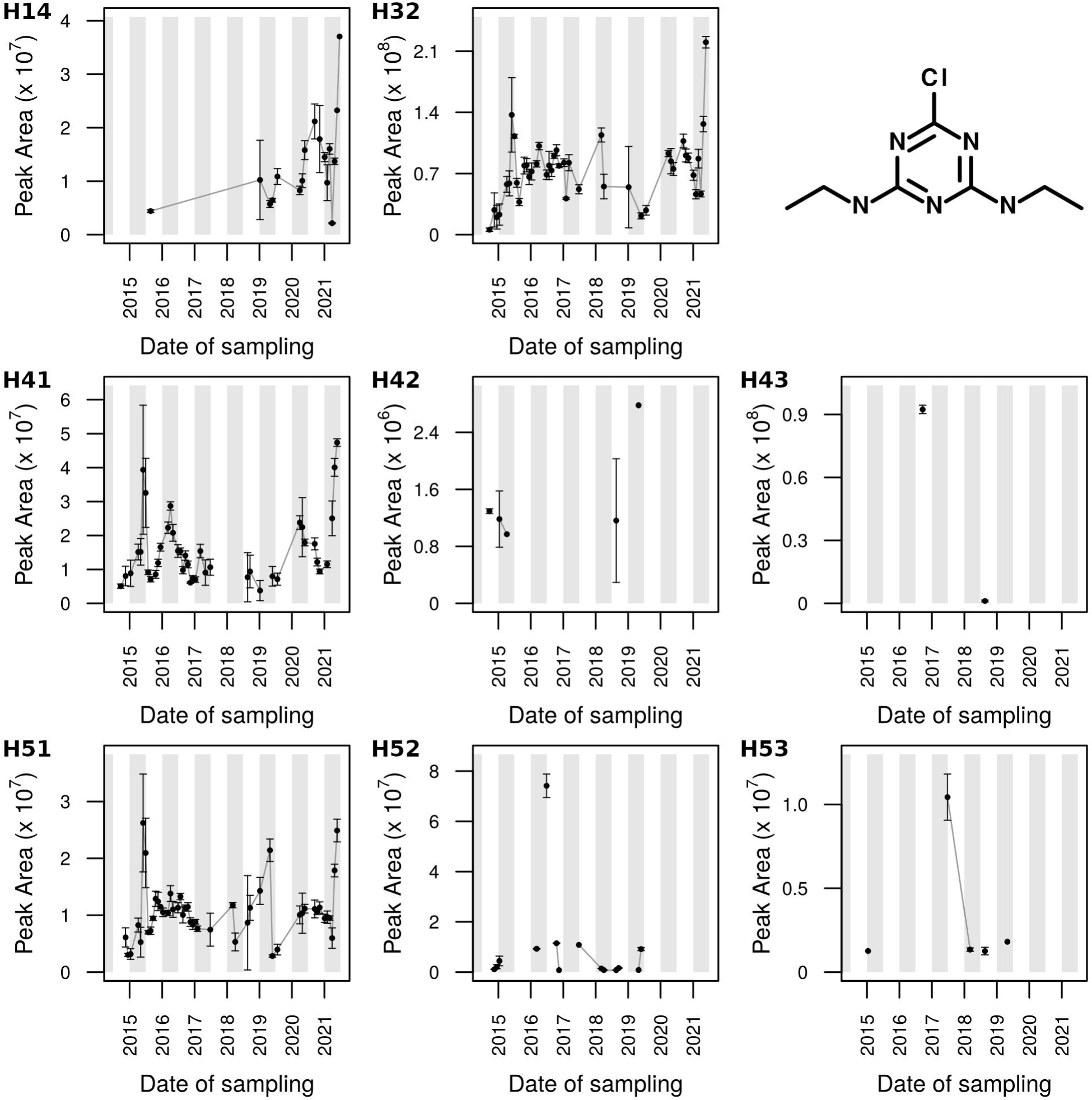

Notably, an increased simazine intensity in the first half of 2021 emerged consistently in perched groundwater (H14 and H32) and in phreatic groundwater of the surface-connected (Lehmann and Totsche, 2020) wells H41 and H51. This suggests that an extraordinary mobilisation event occurred, that may have been caused by strong, but not exceptional rainfall and recharge in February 2021 (Figure 7).

Another notable feature in the dynamics was observed in spring 2015, where simazine intensities increased in wells H32, H41 and H51. This increase coincided with the seasonal trend breakage observed for 7-ODAA, and thus can be attributed equivalently to the period of extreme cumulative precipitation (compare preceding section on 7-ODAA).

The simazine dynamics in H41 exerted two highs in spring 2015 and 2016, respectively, clearly separated by a well-developed minimum in between. Notably, while the first high coincides with a water stand maximum, the second high preceded the 2016 water stand high, though both highs fall into a recharge period (Lehmann and Totsche, 2020; Zerfaß et al., 2022). To that extent, the annual fluctuation of simazine intensity is in response to seasonal recharge patterns and point to lasting release of the legacy contaminant from soil or deeper vadose zones. However, in H51, where water levels showed a similar fluctuation as in H41, the consecutive highs were not as pronounced – rather, an increase throughout 2015 was succeeded by a plateau and a decrease from the second half of 2016 onwards. In further comparing the two wells, H51 only underwent a progressive increase of simazine from 2018 to 2019 (starting in a recharge and progressing in a recession phase). In 2020, both wells showed increasing simazine intensities coinciding with a recharge phase (Zerfaß et al., 2022). Again, this demonstrates that simazine dynamics in the different observers point to differences in recharge intensities and directions. Moreover, as rising water levels and recession phases represent phases of pronounced versus minimum recharge, migration and transformation also continues, and transient flow patterns cause further DOM dynamics.

### 3.4 Hydroxypropazine: A triazine herbicide degradation product sharing dynamics patterns with simazine

As for simazine, hydroxypropazine showed an increase in the first half of 2021 in the wells H14, H32, H41, and H51 (Figure 5). Further, a spring 2015 increase was observed as also for simazine and 7-ODAA in the wells H32, H41 and H51. Further datapoints with elevations in the period 2017-2020 occurred but could not be substantiated due to the low frequency of detection, thus leaving single datapoints with higher intensity from which no trends can be inferred. The wells of less surface-connected groundwater (H42, H43, and H52, H53) were mostly devoid from hydroxypropazine, similar as for simazine.

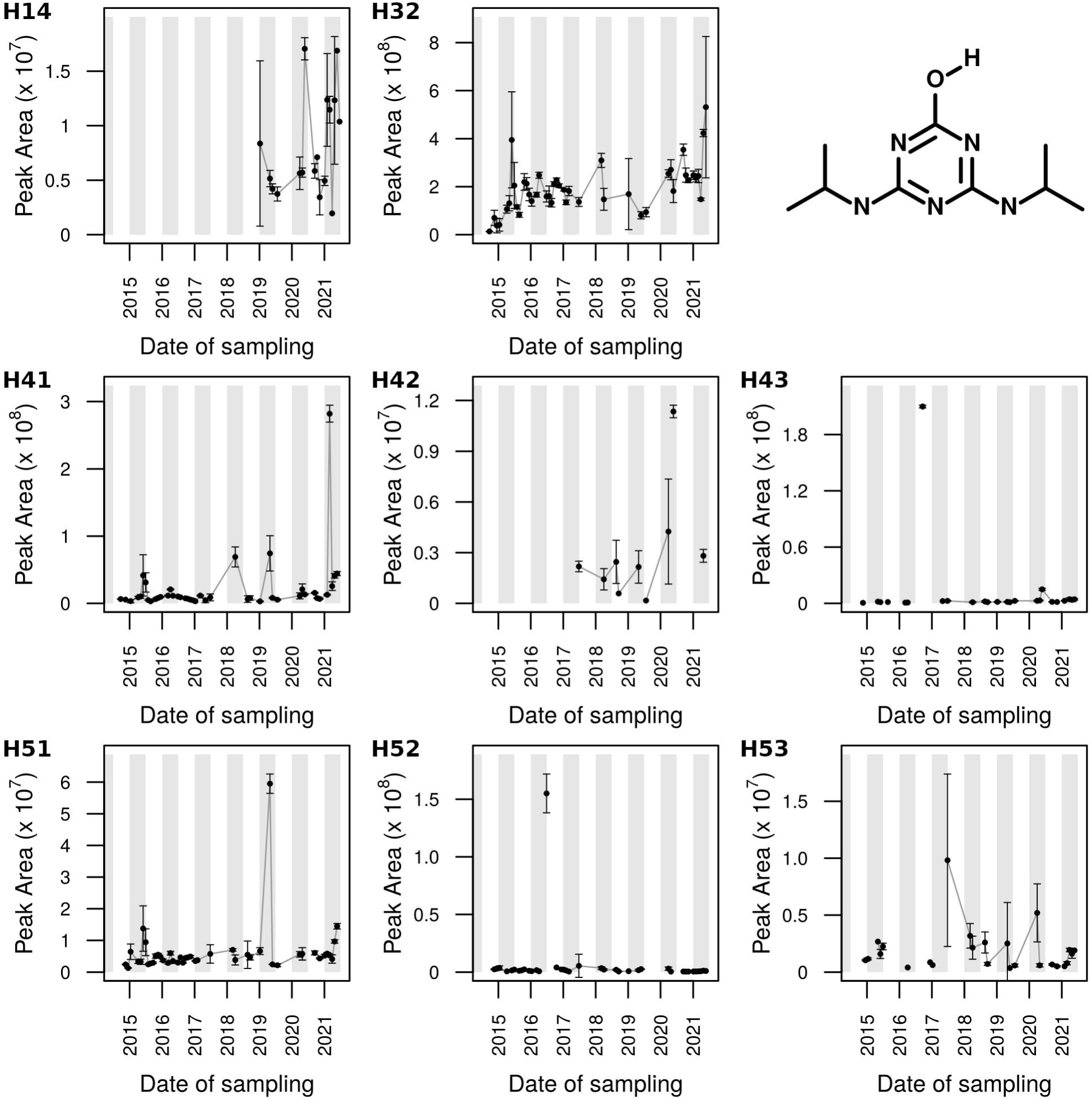

The similarity between the simazine and hydroxypropazine traces suggests that both legacy contaminants likely share application history (parent herbicide for hydroxypropazine) and it is conceivable that they reside in similar soil-related reservoirs with thus coinciding mobilisation dynamics. The presence of hydroxypropazine shows that screening of pesticide metabolites (transformation products) in groundwater is important, as they occasionally were found in higher concentrations than the parent pesticides (Kiefer et al., 2019).

### 3.5 TPP: a concurrent contaminant with severe intensity variability

TPP (Figure 6) remarkably showed severe 2021 intensity increases across all sampling wells, and particularly in the perched groundwater (H14 and H32). For all wells except H51, the 2021 mobilisation also formed the maximum intensity across the entire observation period. H51 only showed a stronger intensity peak in autumn 2015 (consolidated over multiple timepoints of measurement), coinciding with a period of extreme cumulative precipitation at the investigation site (Lehmann and Totsche, 2020). A reason why this may have impacted the TPP dynamics, in contrast to simazine, 7-ODAA and hydroxypropazine, cannot be readily inferred.

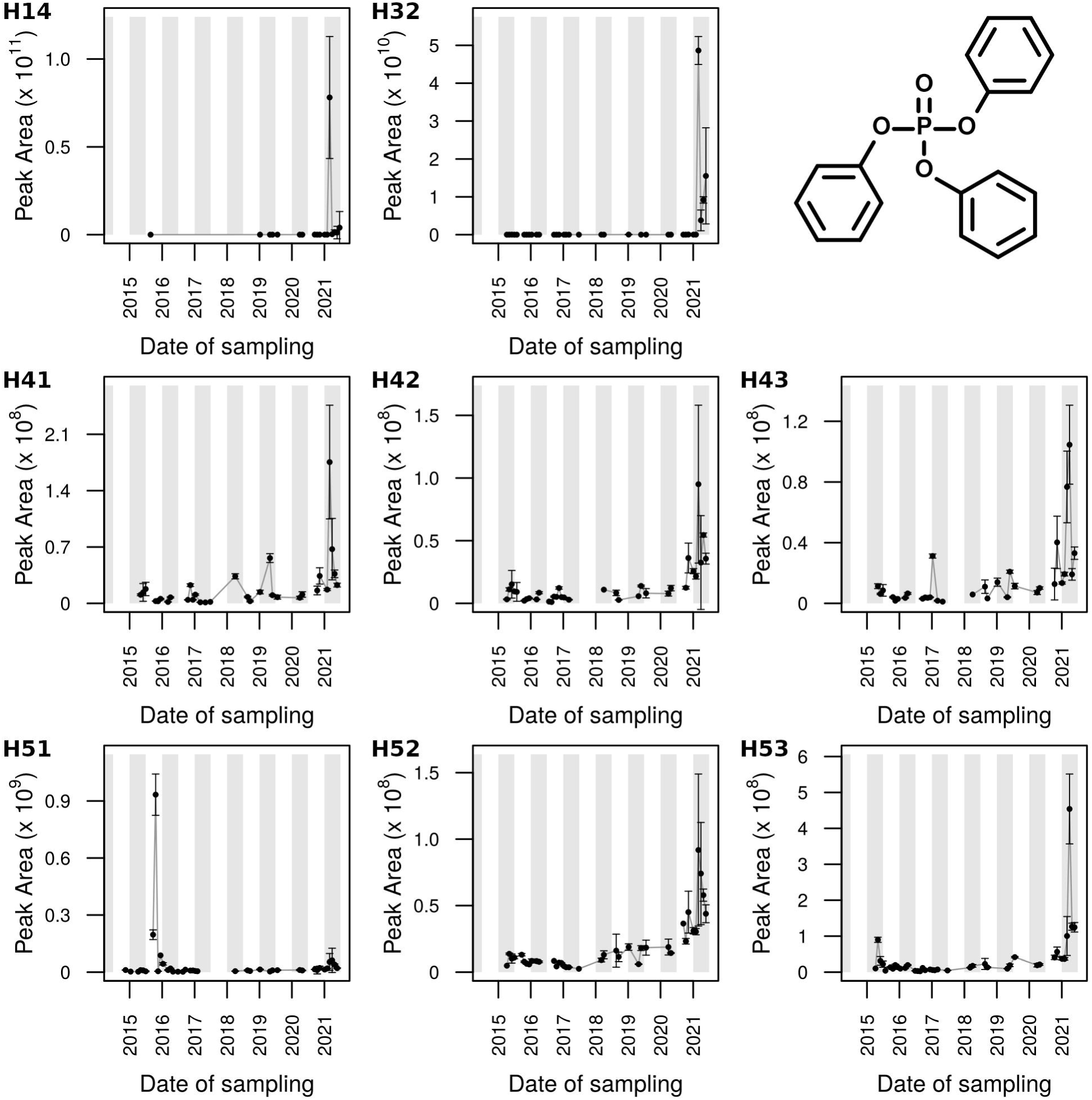

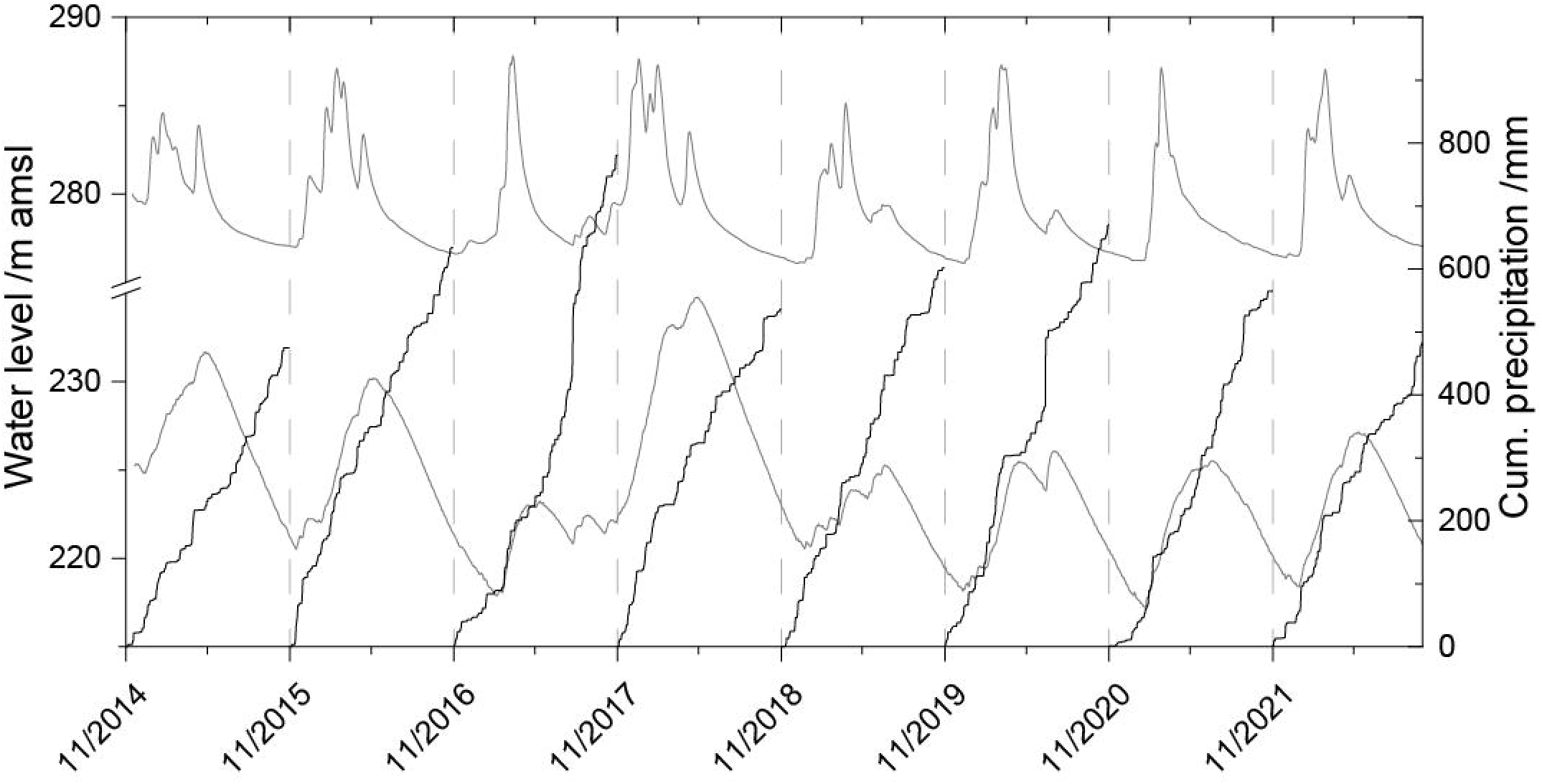

Beyond the 2021 high across the flow system, and the 2015 high in H51, the TPP intensity traces demonstrated the presence of TPP, but no characteristic dynamics, during most of the observation period. As noted, TPP can spread via aerosols, and has a wide use as lubricant additive and flame retardant, so that episodic and diffuse, rather than seasonal release is conceivable. Nevertheless, groundwater loading can occur, as demonstrated by the 2021 dynamics.

In general, untargeted screening is suited to identify contaminations (potential pollutions) that are not included in target screenings (or parameter lists; (Kiefer et al., 2019)), and also allow for discoveries of novel transformation products (Nanusha et al., 2023). Diverse pesticide transformation products, such as hydroxypropazine, can be more soluble and mobile than the parent compounds, making them micropollutants for aquatic ecosystems, which can also have adverse effects on human health (Araya et al., 2024; Nanusha et al., 2023). Moreover, both pesticides and their metabolites can have side effects (other than intended effects) on microbial communities and whole food-webs. Thus, ubiquitous contaminations also threat ecosystem services (e.g., nutrient cycling, degradation of contaminants) in general (Michel et al., 2021).

Furthermore, considering the lag times of regulation measures due to retardation within vadose zone and travel in groundwater, long-term assessments of groundwater quality to pesticide applications are essential for evaluating the effectiveness of regulations (Kim et al., 2022).

## 4. Conclusion

With (ground-)water resources increasingly facing multiple threats of global change (e.g., land-use intensification and climate extremes), in-depth monitoring strategies like untargeted metabolomics monitoring are urgently required to investigate the functioning and stability of groundwater resources. To understand critical zone processes, tracing the in- and outflow of organic matter can shed light on ecosystem functioning and inputs. Our LC-MS investigation identified concurrent and legacy contaminants in groundwater and demonstrates that the dynamics of compounds over the course of years are shaped by multiple controlling factors. Overall, occurrence and intra-seasonal variation of legacy and current-use/release substances both in shallow perched and deep phreatic groundwater (down to ∼90 m) showed local vulnerability to organic contamination in fractured bedrock. Further, the data provides evidence for mobilisation events for both recent, and decades-old legacy contaminants retained in the ground, emerging across the flow system. We note the coincidence of these events with periods of increased water flows or precipitation, representing conceivable impacts to mobilise soil-retained organic matter. The pronounced phases of non-detects, together with cross-compartment observations (e.g. inventory in soil and vadose zone) will further help to investigate surface-subsurface coupling but readily illustrates the need for spatiotemporally high-resolved, non-targeted monitoring. The time traces we present call for further long-term oriented research targeting dissolved organic matter, as obtaining time traces will be paramount to disentangle different influences over time that shape biogeochemical transformations and ecosystem, properties, functions and services. The in part large intensity-fluctuations of the contaminants in groundwater at different timepoints are furthermore a note of caution for regulatory monitoring, as regulatory threshold concentrations until now do not address temporal variability.

## Author Contributions

**Christian Zerfaß:** Conceptualization, Methodology, Formal analysis, Investigation, Data Curation, Writing - Original Draft, Visualization. **Robert Lehmann:** Conceptualization, Methodology, Formal analysis, Investigation, Writing - Review & Editing, Visualization. **Nico Ueberschaar:** Methodology, Investigation, Writing - Review & Editing. **Kai Uwe Totsche:** Conceptualization, Methodology, Investigation, Resources, Writing - Review & Editing, Funding acquisition. **Georg Pohnert:** Conceptualization, Methodology, Investigation, Resources, Writing - Review & Editing, Funding acquisition.

## Acknowledgements

This study is part of the Collaborative Research Centre AquaDiva of the Friedrich Schiller University Jena, funded by the Deutsche Forschungsgemeinschaft (DFG, German Research Foundation) – Project-ID 218627073 – SFB 1076. We thank Felix Kellner for his support in experimentation during his applied laboratory practical internship.

