## Supplementary material for "Current-use and legacy contaminants evidence dissolved organic matter transfer and dynamics across a fractured-rock groundwater recharge area": Figure S2

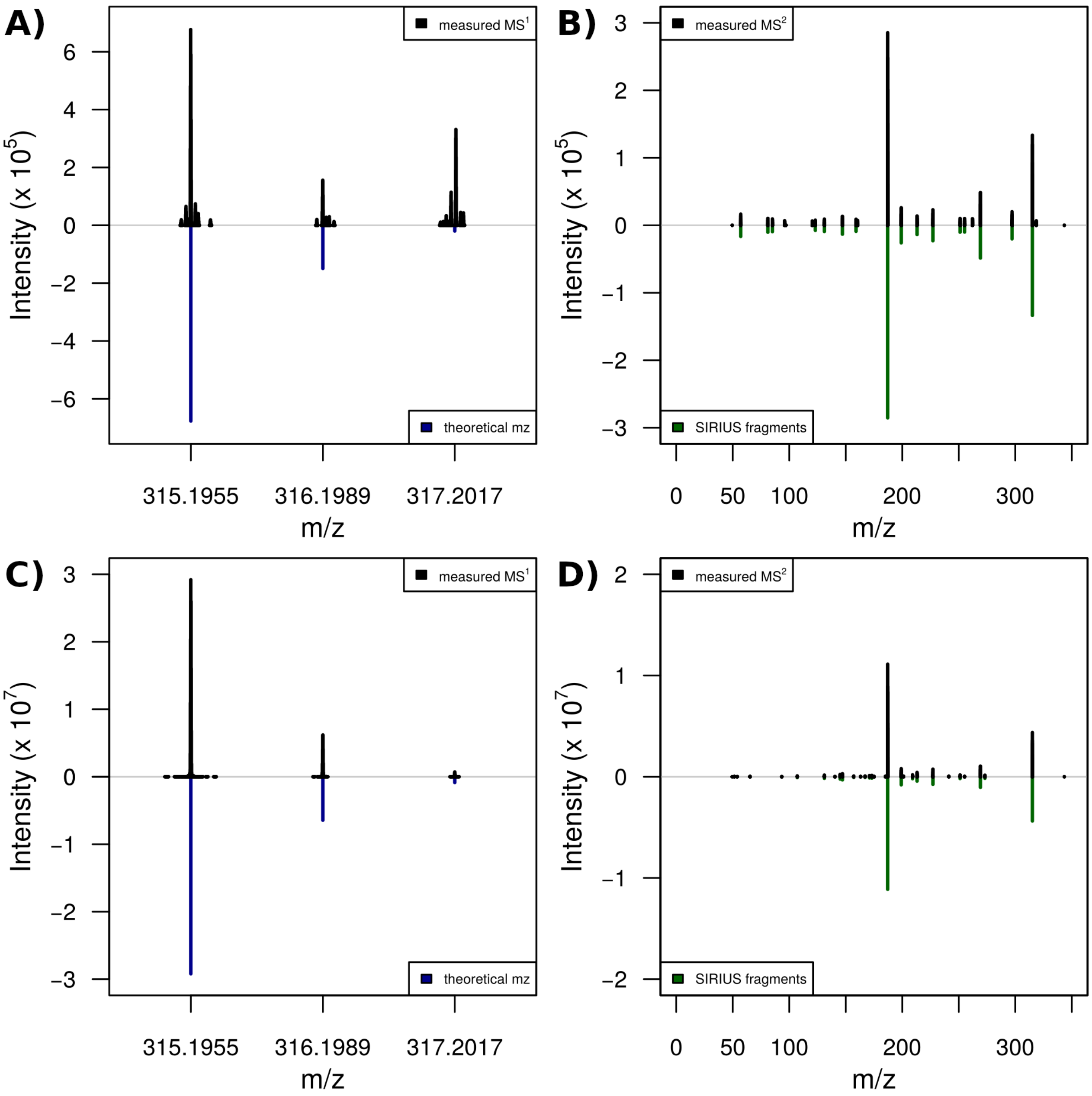


**Supplementary Figure S2.** Identification of 7-ODAA by MS2 ($C_{20}H_{26}O_3$ $[M+H]^+$, monoisotopic theoretical mz = 315.1955) from a groundwater sample (panels A, B, sample taken in February 2021) by comparison with a reference standard (panels C, D). For structure identification, isotope signals in MS1 (A, C) as well as fragments in MS2 (B, D) were taken into consideration. In A and C, measured MS1 signals (black) were compared against theoretical isotope distribution patterns (M, M+1, M+2; blue). In B and D, measured (black) fragment signals and were annotated with suitable fragments by Sirius (green). The retention times of the respective MS2 measurements were 6.7 minutes for the groundwater sample, and 6.8 minutes for the pure reference substance injection. For the MS1 spectra in A, C, the x-axis indicates the mz of the first three isotope signals. Detailed analysis of signals in Supplementary Tables S3, S4.
