## Supplementary material for "Current-use and legacy contaminants evidence dissolved organic matter transfer and dynamics across a fractured-rock groundwater recharge area": Table S1

**Supplementary Table S1.** Detection of DEET by fragmentation analysis (MS2) for a groundwater sample; details to Supplementary Figure S1. The table shows fragments assigned by Sirius. Fragments matching the assignment for the pure reference substance (compare: Supplementary Table S2) are shown in bold.

| **mz signal** | **rel. int.** | **formula** | **adduct** | **exact mass** | **Δppm** |
|---|---|---|---|---|---|
| **72.0442** | **1.5** | **$C_3H_5NO$** | **$[M+H]^+$** | **72.0444** | **-2.8** |
| 72.0808 | 0.6 | $C_4H_9N$ | $[M+H]^+$ | 72.0808 | 0.0 |
| **91.0533** | **0.6** | **$C_7H_6$** | **$[M+H]^+$** | **91.0542** | **-9.9** |
| **100.0748** | **5.2** | **$C_5H_9NO$** | **$[M+H]^+$** | **100.0757** | **-9.0** |
| **119.0486** | **67.8** | **$C_8H_6O$** | **$[M+H]^+$** | **119.0491** | **-4.2** |
| **192.1375** | **100.0** | **$C_{12}H_{17}NO$** | **$[M+H]^+$** | **192.1383** | **-4.2** |
