## Supplementary material for "Current-use and legacy contaminants evidence dissolved organic matter transfer and dynamics across a fractured-rock groundwater recharge area": Table S2

**Supplementary Table S2.** Verification of DEET by fragmentation analysis (MS2) for a pure reference substance injection; details to Supplementary Figure S1. The table shows fragments assigned by Sirius. Fragments matching the assignment for the groundwater sample (compare: Supplementary Table S1) are shown in bold.

| mz signal | rel. int. | formula | adduct | exact mass | Δppm |
|---|---|---|---|---|---|
| **72.0451** | **1.5** | **$C_3H_5NO$** | **$[M+H]^+$** | **72.0444** | **9.7** |
| **91.0540** | **0.6** | **$C_7H_6$** | **$[M+H]^+$** | **91.0542** | **-2.2** |
| **100.0765** | **6.2** | **$C_5H_9NO$** | **$[M+H]^+$** | **100.0757** | **8.0** |
| 109.0655 | 0.5 | $C_7H_8O$ | $[M+H]^+$ | 109.0648 | 6.4 |
| **119.0494** | **84.8** | **$C_8H_6O$** | **$[M+H]^+$** | **119.0491** | **2.5** |
| **192.1385** | **100.0** | **$C_{12}H_{17}NO$** | **$[M+H]^+$** | **192.1383** | **1.0** |
