## Supplementary material for "Current-use and legacy contaminants evidence dissolved organic matter transfer and dynamics across a fractured-rock groundwater recharge area": Table S3

**Supplementary Table S3.** Detection of 7-ODAA by fragmentation analysis (MS2) for a groundwater sample; details to Supplementary Figure S2. The table shows fragments assigned by Sirius. Fragments matching the assignment for the pure reference substance (compare: Supplementary Table S4) are shown in bold.

| mz signal | rel. int. | formula | adduct | exact mass | Δppm |
|---|---|---|---|---|---|
| 57.0700 | 5.8 | $C_4H_8$ | $[M+H]^+$ | 57.0699 | 1.8 |
| 81.0699 | 3.6 | $C_6H_8$ | $[M+H]^+$ | 81.0699 | 0 |
| 85.1007 | 3.3 | $C_6H_{12}$ | $[M+H]^+$ | 85.1012 | -5.9 |
| 123.0804 | 2.7 | $C_8H_{10}O$ | $[M+H]^+$ | 123.0804 | 0 |
| **131.0851** | **3.2** | $\mathbf{C_{10}H_{10}}$ | $\mathbf{[M+H]^+}$ | **131.0855** | **-3.1** |
| **147.0809** | **4.6** | $\mathbf{C_{10}H_{10}O}$ | $\mathbf{[M+H]^+}$ | **147.0804** | **3.4** |
| 159.1157 | 3.1 | $C_{12}H_{14}$ | $[M+H]^+$ | 159.1168 | -6.9 |
| **187.1119** | **100** | $\mathbf{C_{13}H_{14}O}$ | $\mathbf{[M+H]^+}$ | **187.1117** | **1.1** |
| **199.1122** | **9.1** | $\mathbf{C_{14}H_{14}O}$ | $\mathbf{[M+H]^+}$ | **199.1117** | **2.5** |
| **213.1260** | **4.8** | $\mathbf{C_{15}H_{16}O}$ | $\mathbf{[M+H]^+}$ | **213.1274** | **-6.6** |
| **227.1423** | **8.1** | $\mathbf{C_{16}H_{18}O}$ | $\mathbf{[M+H]^+}$ | **227.1430** | **-3.1** |
| **251.1787** | **3.6** | $\mathbf{C_{19}H_{22}}$ | $\mathbf{[M+H]^+}$ | **251.1794** | **-2.8** |
| 255.1755 | 3.5 | $C_{18}H_{22}O$ | $[M+H]^+$ | 255.1743 | 4.7 |
| **269.1911** | **17** | $\mathbf{C_{19}H_{24}O}$ | $\mathbf{[M+H]^+}$ | **269.1900** | **4.1** |
| 297.1839 | 7.1 | $C_{20}H_{24}O_2$ | $[M+H]^+$ | 297.1849 | -3.4 |
| **315.1943** | **46.8** | $\mathbf{C_{20}H_{26}O_3}$ | $\mathbf{[M+H]^+}$ | **315.1955** | **-3.8** |
