## Supplementary material for "Current-use and legacy contaminants evidence dissolved organic matter transfer and dynamics across a fractured-rock groundwater recharge area": Table S4

**Supplementary Table S4.** Verification of 7-ODAA by fragmentation analysis (MS2) for a pure reference substance injection; details to Supplementary Figure S2. The table shows fragments assigned by Sirius. Fragments matching the assignment for the groundwater sample (compare: Supplementary Table S3) are shown in bold.

| mz signal | rel. int. | formula | adduct | exact mass | Δppm |
|---|---|---|---|---|---|
| 107.0853 | 0.8 | $C_8H_{10}$ | $[M+H]^+$ | 107.0855 | -1.9 |
| **131.0851** | **1.4** | $\mathbf{C_{10}H_{10}}$ | $\mathbf{[M+H]^+}$ | **131.0855** | **-3.1** |
| 147.0642 | 2.1 | $C_{10}H_8O$ | $[M+H]^+$ | 145.0648 | -4.1 |
| **147.0802** | **2.7** | $\mathbf{C_{10}H_{10}O}$ | $\mathbf{[M+H]^+}$ | **147.0804** | **-1.4** |
| 147.1174 | 0.6 | $C_{11}H_{14}$ | $[M+H]^+$ | 147.1168 | 4.1 |
| 157.0651 | 0.6 | $C_{11}H_8O$ | $[M+H]^+$ | 157.0648 | 1.9 |
| 167.1064 | 0.6 | $C_{10}H_{14}O_2$ | $[M+H]^+$ | 167.1067 | -1.8 |
| 171.0811 | 1.1 | $C_{12}H_{10}O$ | $[M+H]^+$ | 171.0804 | 4.1 |
| 173.0967 | 1.2 | $C_{12}H_{12}O$ | $[M+H]^+$ | 173.0961 | 3.5 |
| **187.1122** | **100.0** | $\mathbf{C_{13}H_{14}O}$ | $\mathbf{[M+H]^+}$ | **187.1117** | **2.7** |
| **199.1120** | **7.2** | $\mathbf{C_{14}H_{14}O}$ | $\mathbf{[M+H]^+}$ | **199.1117** | **1.5** |
| 209.1332 | 1.8 | $C_{16}H_{16}$ | $[M+H]^+$ | 209.1325 | 3.3 |
| **213.1281** | **3.9** | $\mathbf{C_{15}H_{16}O}$ | $\mathbf{[M+H]^+}$ | **213.1274** | **3.3** |
| **227.1444** | **6.8** | $\mathbf{C_{16}H_{18}O}$ | $\mathbf{[M+H]^+}$ | **227.1430** | **6.2** |
| 227.1782 | 1.4 | $C_{17}H_{22}$ | $[M+H]^+$ | 227.1794 | -5.3 |
| **251.1805** | **1.6** | $\mathbf{C_{19}H_{22}}$ | $\mathbf{[M+H]^+}$ | **251.1794** | **4.4** |
| **269.1883** | **9.4** | $\mathbf{C_{19}H_{24}O}$ | $\mathbf{[M+H]^+}$ | **269.1900** | **-6.3** |
| 273.1485 | 1.4 | $C_{17}H_{20}O_3$ | $[M+H]^+$ | 273.1485 | 0.0 |
| **315.1951** | **39.3** | $\mathbf{C_{20}H_{26}O_3}$ | $\mathbf{[M+H]^+}$ | **315.1955** | **-1.3** |
