## Supplementary material for "Current-use and legacy contaminants evidence dissolved organic matter transfer and dynamics across a fractured-rock groundwater recharge area": Table S5

**Supplementary Table S5.** Detection of simazine by fragmentation analysis (MS2) for a groundwater sample; details to Supplementary Figure S3. The table shows fragments assigned by Sirius. Fragments matching the assignment for the pure reference substance (compare: Supplementary Table S6) are shown in bold.

| mz signal | rel. int. | formula | adduct | exact mass | Δppm |
|---|---|---|---|---|---|
| **68.0249** | **0.3** | **$C_2HN_3$** | **$[M+H]^+$** | **68.0243** | **8.8** |
| **71.0611** | **0.5** | **$C_3H_6N_2$** | **$[M+H]^+$** | **71.0604** | **9.9** |
| **96.0552** | **0.8** | **$C_4H_5N_3$** | **$[M+H]^+$** | **96.0556** | **-4.2** |
| **104.0018** | **0.5** | **$C_2H_2ClN_3$** | **$[M+H]^+$** | **104.0010** | **7.7** |
| **124.0875** | **6.7** | **$C_6H_9N_3$** | **$[M+H]^+$** | **124.0869** | **4.8** |
| **132.0322** | **5.4** | **$C_4H_6ClN_3$** | **$[M+H]^+$** | **132.0323** | **-0.8** |
| **166.1086** | **0.7** | **$C_7H_{11}N_5$** | **$[M+H]^+$** | **166.1087** | **-0.6** |
| **174.0548** | **2.0** | **$C_5H_8ClN_5$** | **$[M+H]^+$** | **174.0541** | **4.0** |
| **202.0859** | **100.0** | **$C_7H_{12}ClN_5$** | **$[M+H]^+$** | **202.0854** | **2.5** |
