## Supplementary material for "Current-use and legacy contaminants evidence dissolved organic matter transfer and dynamics across a fractured-rock groundwater recharge area": Table S6

**Supplementary Table S6.** Verification of simazine by fragmentation analysis (MS2) for a pure reference substance injection; details to Supplementary Figure S3. The table shows fragments assigned by Sirius. Fragments matching the assignment for the groundwater sample (compare: Supplementary Table S5) are shown in bold.

| mz signal | rel. int. | formula | adduct | exact mass | Δppm |
|---|---|---|---|---|---|
| 61.9798 | 0.1 | $CClN$ | $[M+H]^+$ | 61.9792 | 9.7 |
| **68.0248** | **0.2** | $\mathbf{C_2HN_3}$ | $\mathbf{[M+H]^+}$ | **68.0243** | **7.4** |
| **71.0607** | **0.6** | $\mathbf{C_3H_6N_2}$ | $\mathbf{[M+H]^+}$ | **71.0604** | **4.2** |
| 79.0060 | 0.1 | $CH_3ClN_2$ | $[M+H]^+$ | 79.0058 | 2.5 |
| 90.0109 | 0.1 | $C_3H_4ClN$ | $[M+H]^+$ | 90.0105 | 4.4 |
| **96.0553** | **1.0** | $\mathbf{C_4H_5N_3}$ | $\mathbf{[M+H]^+}$ | **96.0556** | **-3.1** |
| **104.0015** | **0.8** | $\mathbf{C_2H_2ClN_3}$ | $\mathbf{[M+H]^+}$ | **104.0010** | **4.8** |
| 107.0373 | 0.1 | $C_3H_7ClN_2$ | $[M+H]^+$ | 107.0371 | 1.9 |
| **124.0865** | **9.5** | $\mathbf{C_6H_9N_3}$ | $\mathbf{[M+H]^+}$ | **124.0869** | **-3.2** |
| **132.0324** | **7.9** | $\mathbf{C_4H_6ClN_3}$ | $\mathbf{[M+H]^+}$ | **132.0323** | **0.8** |
| 138.0778 | 0.1 | $C_5H_7N_5$ | $[M+H]^+$ | 138.0774 | 2.9 |
| **166.1088** | **1.2** | $\mathbf{C_7H_{11}N_5}$ | $\mathbf{[M+H]^+}$ | **166.1087** | **0.6** |
| **174.0547** | **2.5** | $\mathbf{C_5H_8ClN_5}$ | $\mathbf{[M+H]^+}$ | **174.0541** | **3.4** |
| 187.0628 | 0.1 | $C_6H_9ClN_5$ | $[M+H]^+$ | 187.0619 | 4.8 |
| **202.0853** | **100.0** | $\mathbf{C_7H_{12}ClN_5}$ | $\mathbf{[M+H]^+}$ | **202.0854** | **-0.5** |
