## Supplementary material for "Current-use and legacy contaminants evidence dissolved organic matter transfer and dynamics across a fractured-rock groundwater recharge area": Table S7

**Supplementary Table S7.** Detection of hydroxypropazine by fragmentation analysis (MS2) for a groundwater sample; details to Supplementary Figure S4. The table shows fragments assigned by Sirius. Fragments matching the assignment for the pure reference substance (compare: Supplementary Table S8) are shown in bold.

| mz signal | rel. int. | formula | adduct | exact mass | Δppm |
|---|---|---|---|---|---|
| **86.0350** | **0.6** | **$C_2H_3N_3O$** | **$[M+H]^+$** | **86.0349** | **1.2** |
| 86.0600 | 0.3 | $C_2H_5N_4$ | $[M+H]^+$ | 86.0587 | 15.1 |
| 87.0438 | 0.4 | $C_2H_4N_3O$ | $[M+H]^+$ | 87.0427 | 12.6 |
| **94.0651** | **1.6** | **$C_6H_7N$** | **$[M+H]^+$** | **49.0651** | **0.0** |
| 98.0607 | 0.5 | $C_3H_5N_4$ | $[M+H]^+$ | 98.0587 | 20.4 |
| 107.0860 | 0.3 | $C_8H_{10}$ | $[M+H]^+$ | 107.0855 | 4.7 |
| 110.0604 | 0.2 | $C_6H_7NO$ | $[M+H]^+$ | 110.0600 | 3.6 |
| 119.0614 | 1.2 | $C_7H_6N_2$ | $[M+H]^+$ | 119.0604 | 8.4 |
| 121.1018 | 0.2 | $C_9H_{12}$ | $[M+H]^+$ | 121.1012 | 5.0 |
| 124.0756 | 0.7 | $C_5H_7N_4$ | $[M+H]^+$ | 124.0743 | 10.5 |
| 124.1123 | 0.3 | $C_8H_{13}N$ | $[M+H]^+$ | 124.1121 | 1.6 |
| 126.0549 | 0.2 | $C_4H_5N_4O$ | $[M+H]^+$ | 126.0536 | 10.3 |
| 126.0914 | 2.3 | $C_7H_{11}NO$ | $[M+H]^+$ | 126.0913 | 0.8 |
| 127.0746 | 0.4 | $C_5H_8N_3O$ | $[M+H]^+$ | 127.0740 | 4.7 |
| **128.0547** | **5.5** | **$C_3H_5N_5O$** | **$[M+H]^+$** | **128.0567** | **-5.5** |
| **128.0820** | **1.7** | **$C_5H_9N_3O$** | **$[M+H]^+$** | **128.0818** | **1.6** |
| 138.0914 | 0.4 | $C_6H_9N_4$ | $[M+H]^+$ | 138.0900 | 10.1 |
| 139.0759 | 1.0 | $C_6H_8N_3O$ | $[M+H]^+$ | 139.0740 | 13.7 |
| 150.0916 | 1.1 | $C_7H_9N_4$ | $[M+H]^+$ | 150.0900 | 10.7 |
| 152.1067 | 0.3 | $C_9H_{13}NO$ | $[M+H]^+$ | 152.1070 | -2.0 |

| | | | | | |
|---|---|---|---|---|---|
| 156.1020 | 0.3 | $C_6H_{11}N_4O$ | $[M+H]^+$ | 156.1006 | 9.0 |
| 166.1228 | 0.4 | $C_8H_{13}N_4$ | $[M+H]^+$ | 166.1213 | 9.0 |
| 167.1063 | 0.8 | $C_8H_{12}N_3O$ | $[M+H]^+$ | 167.1053 | 6.0 |
| **170.1049** | **16.6** | **$C_6H_{11}N_5O$** | **$[M+H]^+$** | **170.1036** | **7.6** |
| 176.1072 | 0.3 | $C_9H_{11}N_4$ | $[M+H]^+$ | 176.1056 | 9.1 |
| 194.1168 | 1.6 | $C_9H_{13}N_4O$ | $[M+H]^+$ | 194.1162 | 3.1 |
| **212.1513** | **100.0** | **$C_9H_{17}N_5O$** | **$[M+H]^+$** | **212.1506** | **3.3** |
