## Supplementary material for "Current-use and legacy contaminants evidence dissolved organic matter transfer and dynamics across a fractured-rock groundwater recharge area": Table S8

**Supplementary Table S8.** Verification of hydroxypropazine by fragmentation analysis (MS2) for a pure reference substance injection; details to Supplementary Figure S4. The table shows fragments assigned by Sirius. Fragments matching the assignment for the groundwater sample (compare: Supplementary Table S7) are shown in bold.

| mz signal | rel. int. | formula | adduct | exact mass | Δppm |
|---|---|---|---|---|---|
| **86.0352** | **0.8** | **$C_2H_3N_3O$** | **$[M+H]^+$** | **86.0349** | **3.5** |
| **94.0651** | **1.1** | **$C_6H_7N$** | **$[M+H]^+$** | **94.0651** | **0.0** |
| **128.0562** | **5.1** | **$C_3H_5N_5O$** | **$[M+H]^+$** | **128.0567** | **-3.9** |
| **128.0824** | **1.5** | **$C_5H_9N_3O$** | **$[M+H]^+$** | **128.0818** | **4.7** |
| **170.1044** | **15.6** | **$C_6H_{11}N_5O$** | **$[M+H]^+$** | **170.1036** | **4.7** |
| **212.1507** | **100.0** | **$C_9H_{17}N_5O$** | **$[M+H]^+$** | **212.1506** | **0.5** |
