## Supplementary material for "Current-use and legacy contaminants evidence dissolved organic matter transfer and dynamics across a fractured-rock groundwater recharge area": Table S9

**Supplementary Table S9.** Detection of TPP by fragmentation analysis (MS2) for a groundwater sample; details to Supplementary Figure S5. The table shows fragments assigned by Sirius. Fragments matching the assignment for the pure reference substance (compare: Supplementary Table S10) are shown in bold.

| mz signal | rel. int. | formula | adduct | exact mass | Δppm |
|---|---|---|---|---|---|
| **53.0392** | **0.2** | **$C_4H_4$** | **$[M+H]^+$** | **53.0386** | **11.3** |
| **77.0395** | **0.3** | **$C_6H_4$** | **$[M+H]^+$** | **77.0386** | **11.7** |
| **95.0489** | **1.1** | **$C_6H_6O$** | **$[M+H]^+$** | **95.0491** | **-2.1** |
| 149.0605 | 0.2 | $C_9H_8O_2$ | $[M+H]^+$ | 149.0597 | 5.4 |
| **152.0618** | **1.6** | **$C_{12}H_7$** | **$[M+H]^+$** | **152.0621** | **-2.0** |
| **153.0705** | **6.2** | **$C_{12}H_8$** | **$[M+H]^+$** | **153.0699** | **3.9** |
| **169.0656** | **0.2** | **$C_{12}H_8O$** | **$[M+H]^+$** | **169.0648** | **4.7** |
| **171.0810** | **1.6** | **$C_{12}H_{10}O$** | **$[M+H]^+$** | **171.0804** | **3.5** |
| 175.0190 | 0.3 | $C_{13}H_2O$ | $[M+H]^+$ | 175.0178 | 6.9 |
| **215.0260** | **0.7** | **$C_{12}H_7O_2P$** | **$[M+H]^+$** | **215.0256** | **1.9** |
| **229.1022** | **2.9** | **$C_{18}H_{12}$** | **$[M+H]^+$** | **229.1012** | **4.4** |
| 233.0234 | 0.6 | $C_{15}H_4O_3$ | $[M+H]^+$ | 233.0233 | 0.4 |
| **233.0369** | **11.4** | **$C_{12}H_9O_3P$** | **$[M+H]^+$** | **233.0362** | **3.0** |
| 251.0335 | 0.3 | $C_{15}H_6O_4$ | $[M+H]^+$ | 251.0339 | -1.6 |
| **251.0486** | **4.4** | **$C_{12}H_{11}O_4P$** | **$[M+H]^+$** | **251.0468** | **7.2** |
| **251.0637** | **0.1** | **$C_{16}H_{11}OP$** | **$[M+H]^+$** | **251.0620** | **6.8** |
| **309.0671** | **1.0** | **$C_{18}H_{13}O_3P$** | **$[M+H]^+$** | **309.0675** | **-1.3** |
| **327.0782** | **100.0** | **$C_{18}H_{15}O_4P$** | **$[M+H]^+$** | **327.0781** | **0.3** |
