## Supplementary material for "Current-use and legacy contaminants evidence dissolved organic matter transfer and dynamics across a fractured-rock groundwater recharge area": Table S10

**Supplementary Table S10.** Verification of TPP by fragmentation analysis (MS2) for a pure reference substance injection; details to Supplementary Figure S5. The table shows fragments assigned by Sirius. Fragments matching the assignment for the groundwater sample (compare: Supplementary Table S9) are shown in bold.

| mz signal | rel. int. | formula | adduct | exact mass | Δppm |
|---|---|---|---|---|---|
| **53.0390** | **0.2** | **$C_4H_4$** | **$[M+H]^+$** | **53.0386** | **7.5** |
| **77.0392** | **0.4** | **$C_6H_4$** | **$[M+H]^+$** | **77.0386** | **7.8** |
| **95.0490** | **1.0** | **$C_6H_6O$** | **$[M+H]^+$** | **95.0491** | **-1.1** |
| 98.9844 | 0.1 | $H_3O_4P$ | $[M+H]^+$ | 98.9842 | 2.0 |
| **152.0622** | **1.2** | **$C_{12}H_7$** | **$[M+H]^+$** | **152.0621** | **0.7** |
| **153.0706** | **5.1** | **$C_{12}H_8$** | **$[M+H]^+$** | **153.0699** | **4.6** |
| **169.0635** | **0.2** | **$C_{12}H_8O$** | **$[M+H]^+$** | **169.0648** | **-7.7** |
| **171.0811** | **1.5** | **$C_{12}H_{10}O$** | **$[M+H]^+$** | **171.0804** | **4.1** |
| 175.0145 | 0.4 | $C_6H_7O_4P$ | $[M+H]^+$ | 175.0155 | -5.7 |
| 202.0779 | 0.1 | $C_{16}H_9$ | $[M+H]^+$ | 202.0777 | 1.0 |
| **215.0259** | **0.7** | **$C_{12}H_7O_2P$** | **$[M+H]^+$** | **215.0256** | **1.4** |
| 227.0847 | 0.2 | $C_{18}H_{10}$ | $[M+H]^+$ | 227.0855 | -3.5 |
| 228.0945 | 1.5 | $C_{18}H_{11}$ | $[M+H]^+$ | 228.0934 | 4.8 |
| **229.1009** | **2.4** | **$C_{18}H_{12}$** | **$[M+H]^+$** | **229.1012** | **-1.3** |
| 233.0151 | 0.2 | $C_{15}H_5OP$ | $[M+H]^+$ | 233.0151 | 0.0 |
| **233.0353** | **9.9** | **$C_{12}H_9O_3P$** | **$[M+H]^+$** | **233.0362** | **-3.9** |
| **251.0465** | **4.1** | **$C_{12}H_{11}O_4P$** | **$[M+H]^+$** | **251.0468** | **-1.2** |
| **251.0616** | **0.2** | **$C_{16}H_{11}OP$** | **$[M+H]^+$** | **251.0620** | **4.1** |
| 291.0581 | 0.1 | $C_{18}H_{11}O_2P$ | $[M+H]^+$ | 291.0569 | 4.1 |
| **309.0682** | **0.9** | **$C_{18}H_{13}O_3P$** | **$[M+H]^+$** | **309.0675** | **2.3** |

| | | | | | |
|---|---|---|---|---|---|
| **327.0778** | **100.0** | **$C_{18}H_{15}O_4P$** | **$[M+H]^+$** | **327.0781** | **-0.9** |
